# Unexpected genetic and microbial diversity for arsenic cycling in deep sea cold seep sediments

**DOI:** 10.1101/2022.11.20.517286

**Authors:** Chuwen Zhang, Xinyue Liu, Ling-Dong Shi, Jiwei Li, Xi Xiao, Zongze Shao, Xiyang Dong

## Abstract

Cold seeps, where cold hydrocarbon-rich fluid escapes from the seafloor, showed strong enrichment of toxic metalloid arsenic (As). The toxicity and mobility of As can be greatly altered by microbial processes that play an important role in global As biogeochemical cycling. However, a global overview of genes and microbes involved in As transformation at seeps remains to be fully unveiled. Using 87 sediment metagenomes and 33 metatranscriptomes derived from 13 globally distributed cold seeps, we show that As detoxification genes (*arsM, arsP, arsC1*/*arsC*2, *acr3*) were prevalent at seeps and more phylogenetically diverse than previously expected. Asgardarchaeota and a variety of unidentified bacterial phyla (e.g. 4484-113, AABM5-125-24 and RBG-13-66-14) may also function as the key players in As transformation. The abundances of As-cycling genes and the compositions of As-associated microbiome shifted across different sediment depths or types of cold seep. The energy-conserving arsenate reduction or arsenite oxidation could impact biogeochemical cycling of carbon and nitrogen, via supporting carbon fixation, hydrocarbon degradation and nitrogen fixation. Overall, this study provides a comprehensive overview of As-cycling genes and microbes at As-enriched cold seeps, laying a solid foundation for further studies of As cycling in deep sea microbiome at the enzymatic and processual levels.

## Introduction

Cold seeps are characterized by the emission of subsurface fluids into the sea floor and occur widely at active and passive continental margins^1, 2^. The upward fluids are often rich in methane and other hydrocarbons which sustain sea bed oasis composed of various microorganisms and faunal assemblages^3, 4^. The primary process that fuel complex cold seep ecosystems is the anaerobic oxidation of methane (AOM), conjointly operated by a consortium of anaerobic methane-oxidizing archaea (ANME) and sulfate-reducing bacteria (SRB)^5, 6^. AOM removes approximately 80% of upward venting methane, acting as an efficient methane filter^7^. Additionally, deep-sea cold seep sediments also contain diverse and abundant diazotrophs that might contribute substantially to the global nitrogen balance^8^. Cold seeps are therefore biologically and geochemically significant on a global scale.

The venting fluids can significantly influence the sedimentary environment of seep sites, resulting in changes of chemical characteristics of sediments^9^. In particular, arsenic (As), one of the most abundant elements in the Earth’s crust, are anomalously enriched in seep sediments^10-14^. The anomalous As enrichment could be attributed to the ascending fluids that could capture As and other metals when passing through thick shaly formations^10, 14^; or the so-called particulate iron shuttle effect^9, 11, 13^. As is also toxic metalloid in nature that, upon exposure, can cause negative effects for all living things^15^. Depending on the physicochemical conditions, As can be found in different oxidation and methylation states, showing various levels of toxicity and bioavailability^16^. In marine environments, arsenate (As(V)), and arsenite (As(III)) are the dominant forms of inorganic As^17^. It is assumed that microbes have evolved a genetic repertoire related to As cycling, dated back to at least 2.72 billion years ago^18,19^. As biotransformation processes include As detoxification to mitigate toxicity and As respiration to conserve energy. The As detoxification is mainly achieved by two steps: reduction of As(V) to As(III) by cytoplasmic As(V) reductases (*arsC* gene) with homology to either glutaredoxin (*arsC1* gene) or thioredoxin (*arsC2* gene) family and subsequent extrusion of As(III) via As(III) efflux permeases (*arsB* and *acr3* genes)^20,21^ (**Figure 1**). Another As detoxification mechanism involves the methylation of As(III) to methylarsenite (MAs(III)) by the As(III) S-adenosylmethionine (SAM) methyltransferase (*arsM* gene)^22^ (**Figure 1**). Although MAs(III) intermediates are more toxic than As(III), they do not accumulate in cells and can be detoxified through several different pathways. MAs(III) can be further methylated by ArsM and volatilized, extruded from cells via the MAs(III) efflux permease (*arsP* gene)^23^, oxidated to less toxic MAs(V) by the MAs(III)-specific oxidase (*arsH* gene)^24^, or demethylated to less toxic As(III) by the C-As lyase (*arsI* gene)^25^. As respiration consists of the chemolithotrophic oxidation of As(III) by As(III) oxidase (*aioAB*/*arxAB* gene cluster) and dissimilatory As(V) reduction by respiratory As(V) reductase (*arrAB* gene cluster)^15, 26^ (**Figure 1**). Taken together, microbes have a huge potential effect on the biogeochemical cycling and toxicity of As.

**Figure 1.**
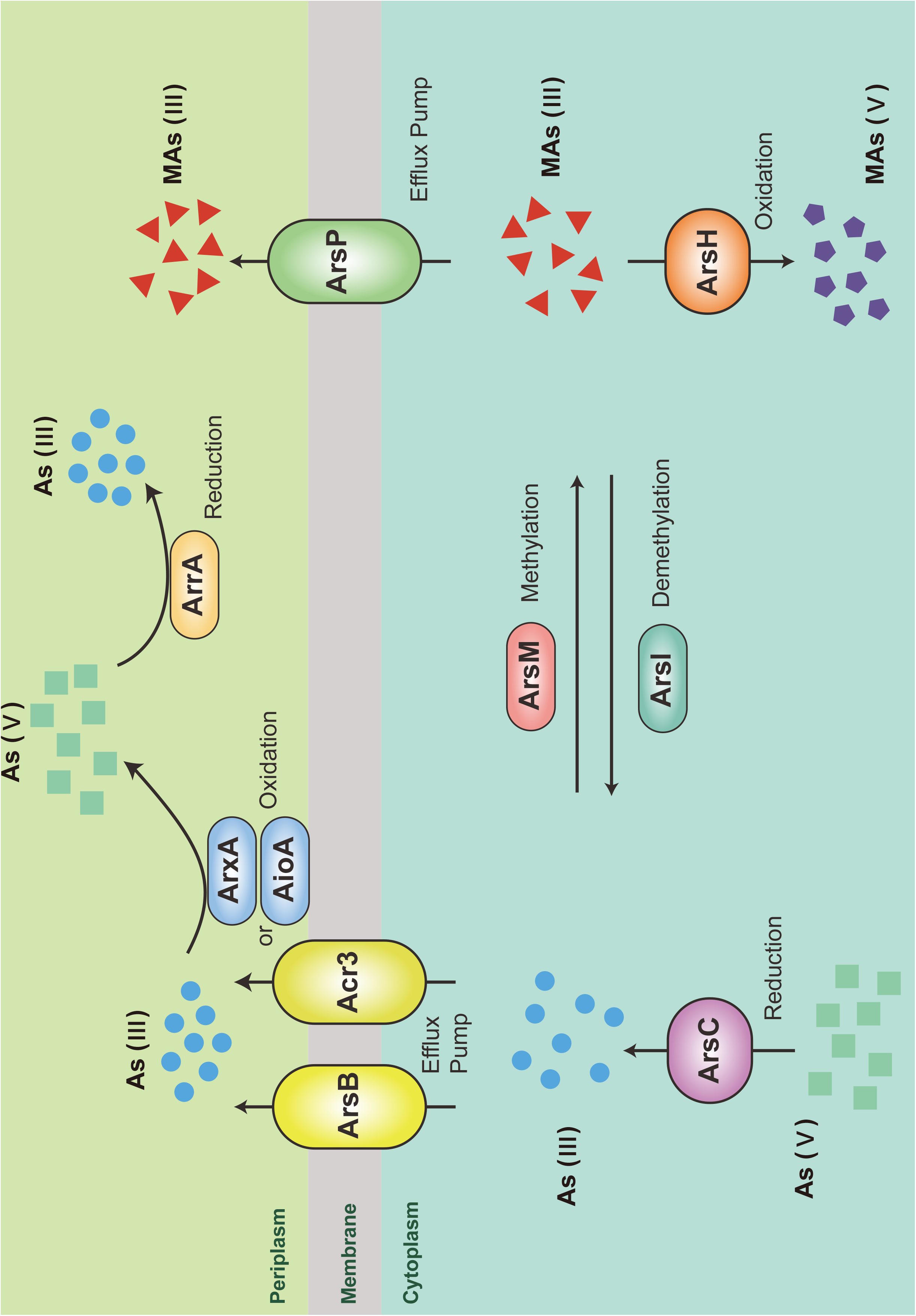
Diagram of the microbial transformations of As. As(III), arsenite; As(V), arsenate; MMAs(III), trivalent methylarsenite; MMAs(V), pentavalentmethylarsenate. As(III) efflux permease: ArsB/Acr3; cytoplasmic As(V) reductase: ArsC; respiratory As(V) reductase: ArrA; As(III) oxidase: AioA/ArxA; As(III) S-adenosylmethionine (SAM) methyltransferase: ArsM; C-As lyase: ArsI; MAs(III) efflux permease: ArsP; MAs(III)-specific oxidase: ArsH.

So far, As-transforming microbes and As-related genes have been widely investigated in various natural environments, including polluted and pristine soils^27, 28^, terrestrial geothermal springs^29-31^, wetlands^32, 33^, pelagic oxygen-deficient zones^34^, groundwater^35^, etc. For example, metagenomic and metatranscriptomic analyses revealed that Aquificae are the key players for the *arsC*-based detoxification in Tengchong geothermal springs^29^. A global survey also described the phylogenetic diversity, genomic location, and biogeography of As-related genes in soil metagenomes^36^. Only recently, the behavior of As biotransformation has been reported in the deep-sea realms, i.e. hadal trench of the Challenger Deep^37^. Deep-sea ecosystems cover 67% of Earth surface and have extremely high densities of microbes (up to 1000× greater than surface waters) which play a critical role for long-term controls on global biogeochemical cycles^38, 39^. The environmental conditions in deep seafloor cold seeps differ greatly from those in the aforementioned ecosystems, such as low temperatures, high pressure, darkness and the presence of seepage activities^1^. Thus, the investigation of As-related genes and microbes at seeps will expand our current knowledge on As metabolisms and allow us to discover new lineages containing As-related genes.

The purpose of this study was to decipher the microbial transformation of As in cold seep sediments at a global scale. Here, we applied a relatively comprehensive data set of 87 sediment metagenomes and 33 metatranscriptomes derived from 13 geographically diverse cold seeps across global oceans (**Supplementary Data 1**), to investigate As-associated genes and their host microbes. This study aims to address the following questions: (i) biogeography of As-cycling genes across global cold seeps; (ii) phylogenetic diversity and distribution of As-cycling genes across global cold seeps; (iii) interactions between As metabolisms and biogeochemical cycles of carbon and nitrogen.

## Results and discussion

### As cycling genes are widespread and active across global cold seeps

To gain a broad view on biogeography of As cycling genes, we determined their abundance from 87 sediment metagenomes collected from 13 globally distributed cold seeps. Considering that sulfate respiration is one of the most important microbial redox processes in cold seep sediments^40^, the *dsrA* was used as the target gene to compare with As cycling genes. We found that genes related to As detoxification were prevalent in these cold seep samples (**Figure 2**) and their abundances were higher than those of *dsrA* genes (**Supplementary Figures 2-3; Supplementary Data 2**). The *arsM* and *arsP* genes that respectively produce volatilized methylated organoarsenicals and mediate its subsequent expulsion outside cell, were the most abundant ones. The *arsC1*/*arsC2* genes for cytoplasmic As(V) reduction and the *acr3* gene for As(III) extrusion also dominated most cold seep samples. Moreover, As detoxification genes, i.e. *arsM, arsP, arsC1*/*arsC2, acr3*, were actively expressed in the sediment metatranscriptomes from Haima and Jiaolong seeps along with gas hydrate deposit zones of Qiongdongnan and Shenhu, revealing *in situ* microbial activities on As detoxification (**Supplementary Figure 4**). Seep microbes might utilize both methylation and cytoplasmic As(V) reduction strategies to overcome potential toxic effects of exceptional As accumulation at cold seeps. Alternatively, methylation is not strictly a detoxification pathway but also an antibiotic-producing process with MAs(III) being a primitive antibiotic^41^, which could provide additional competitive advantages. However, the function of *arsM* in anoxic environments and its contribution to As cycling have yet to be verified. Our results contradict previous findings demonstrating that *arsM* are less common in soils^36^ and hot springs^29^ than *arsC*, but in line with those found in hadal sediments^37^. The discrepancy in As detoxification mechanisms between terrestrial and deep-sea ecosystems could be attributed to their huge variations in the habitats and geographical locations. When comparing the abundance of As(III) efflux pumps, we observed that *arsB* was in much lower abundance than *acr3* (**Figure 2a**). Previous studies also reported an abundance of *acr3* over *arsB* in forest soils and wetlands^32, 36, 42^. This is likely because Acr3 proteins are more ancient and have greater phylogenetic distribution as compared with ArsB^19^. Conversely, genes related to energetic As respiratory oxidation (*aioA*) and reduction (*arrA*/*arxA*) were less abundant in all cold seep samples as compared with As detoxification genes. Despite of this, respiratory genes were transcriptionally active, as evidenced by the detection of *arrA* transcripts in Jiaolong seep (up to 15.9 TPM, **Supplementary Figure 4**).

**Figure 2.**
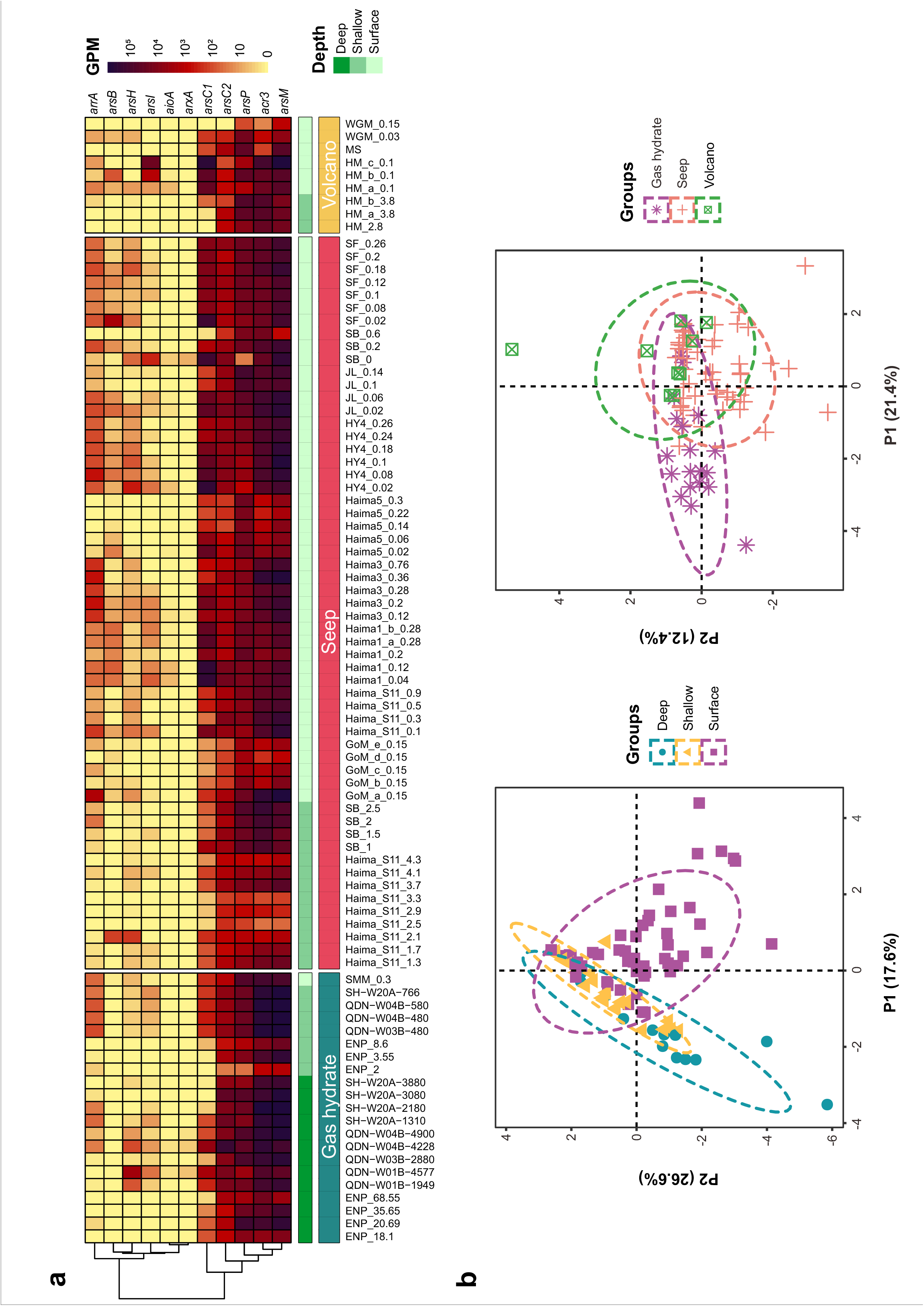
The global distribution of potential genes involved in As cycling at cold seeps. (a) The abundances of As cycling genes across the 87 cold seep metagenomes. The abundance of each gene was normalized by the gene length and sequencing depth and represented as GPM (genes per million) value. (b) The partial least squares discrimination analysis (PLS-DA) plots based on the abundances of As-cycling genes (*n* = 87). Similarity values among the samples of different sediment depths and types of cold seep were examined using a 999-permutation PERMANOVA test. Source data is available in **Supplementary Data 2**.

To determine the distribution characteristics of As cycling genes, each metagenome was categorized in terms of its sediment depth (i.e. surface: <1 mbsf; shallow: 1-10 mbsf; deep: >10 mbsf). Metagenomes were also grouped based on the type of cold seep, including gas hydrate, seep (i.e. oil and gas/methane seep), and volcano (mud/asphalt volcano)^1^, respectively. The partial least squares discrimination analysis (PLS-DA) revealed dissimilarity in As cycling genes among different sediment layers (**Figure 2b**; *F* = 4.3504, *p* = 0.001, *R*^*2*^ = 0.10267, 999-permutations PERMANOVA test). The distribution traits of As cycling genes in surface sediments were separated from deep sediments and more similar to those in shallow ones (**Figure 2b**). The abundance of prevalent As cycling genes such as *acr3, arsC2* and *arsM* in deep sediments were significantly higher as compared with those in shallow and surface sediments (**Supplementary Figure 2**). As cycling genes in different types of cold seep were also different from each other (**Figure 2b**; *F* = 3.5246, *p* = 0.004, *R*^*2*^ = 0.07742, 999-permutations PERMANOVA test). Dominant As cycling genes in gas hydrates displayed higher abundances relative to those in seeps and volcanos (**Supplementary Figure 3**). Hence, the distributions of As-associated genes were influenced by a combination of sediment depths and types of cold seep. The higher As-cycling gene abundance observed in our deep or gas hydrate-associated samples could be correlated with a high level of environmental As, as what described in As-rich altiplanic wetland^32^. In the Nankai Trough, As with unknown sources was demonstrated to actively release into sediments layers where methane hydrates occur (As concentration of 14 ppm in gas hydrate-bearing sediments vs av. 6.4 ppm for the whole sediment core)^17^.

### Microbes involved in As cycling varied between seep habitats

To profile taxonomic diversity of As-related microbes, a total of 1741 species-level metagenome-assembled genomes (MAGs, 95% average nucleotide identity) were reconstructed from these 87 cold seep metagenomes (**Supplementary Data 3**). Of these, 1083 MAGs spanning 9 archaeal and 63 bacterial phyla as well as one unclassified bacterial phylum were potentially involved in As cycling at cold seeps (**Supplementary Data 4**). Metagenomic read recruitments revealed that the recovered 1083 As-related MAGs accounted for 1.8-62.8% cold seep communities (**Figure 3 and Supplementary Data 4**). Taxonomic compositions of As-related microbiomes across different types of cold seep displayed pronounced variations (**Figure 3**). In the sediments derived from oil and gas/methane seep, As-related microbes contained mostly Methanogasteraceae (i.e. ANME-2c) and Methanocomedenaceae (i.e. ANME-2a) within Halobacteriota phylum, ETH-SRB1 within Desulfobacterota phylum, JS1 within Atribacterota phylum as well as Anaerolineae and Dehalococcoidia within Chloroflexota phylum. The As-related microbes in gas hydrate sediments were dominated by bacterial lineages, highlighted by Atribacterota (JS1) and Chloroflexota (Anaerolineae and Dehalococcoidia). Nevertheless, in asphalt/mud volcano sediments, the compositions of As-related microbes were diverse in different samples. The clear distinctions in As-related microbiomes between different seep habitats suggested an important role driven by environment selection. Multiple parameters, including sediment temperature, sediment depth, water depth, methane concentration, and geographic distance have been demonstrated to cause these variations^43, 44^. Additionally, our results show that Chloroflexota outnumbered Atribacterota in sediment samples with lower Fe(II) concentration (av. 16.51 µmol/L), while Atribacterota dominated over Chloroflexota in sediment samples with higher Fe(II) concentration (av. 81.54 µmol/L). It is possible that iron oxyhydroxides control the mobilization of As^45^ and thus affect As-related microbial communities (**Figure 3**).

**Figure 3.**
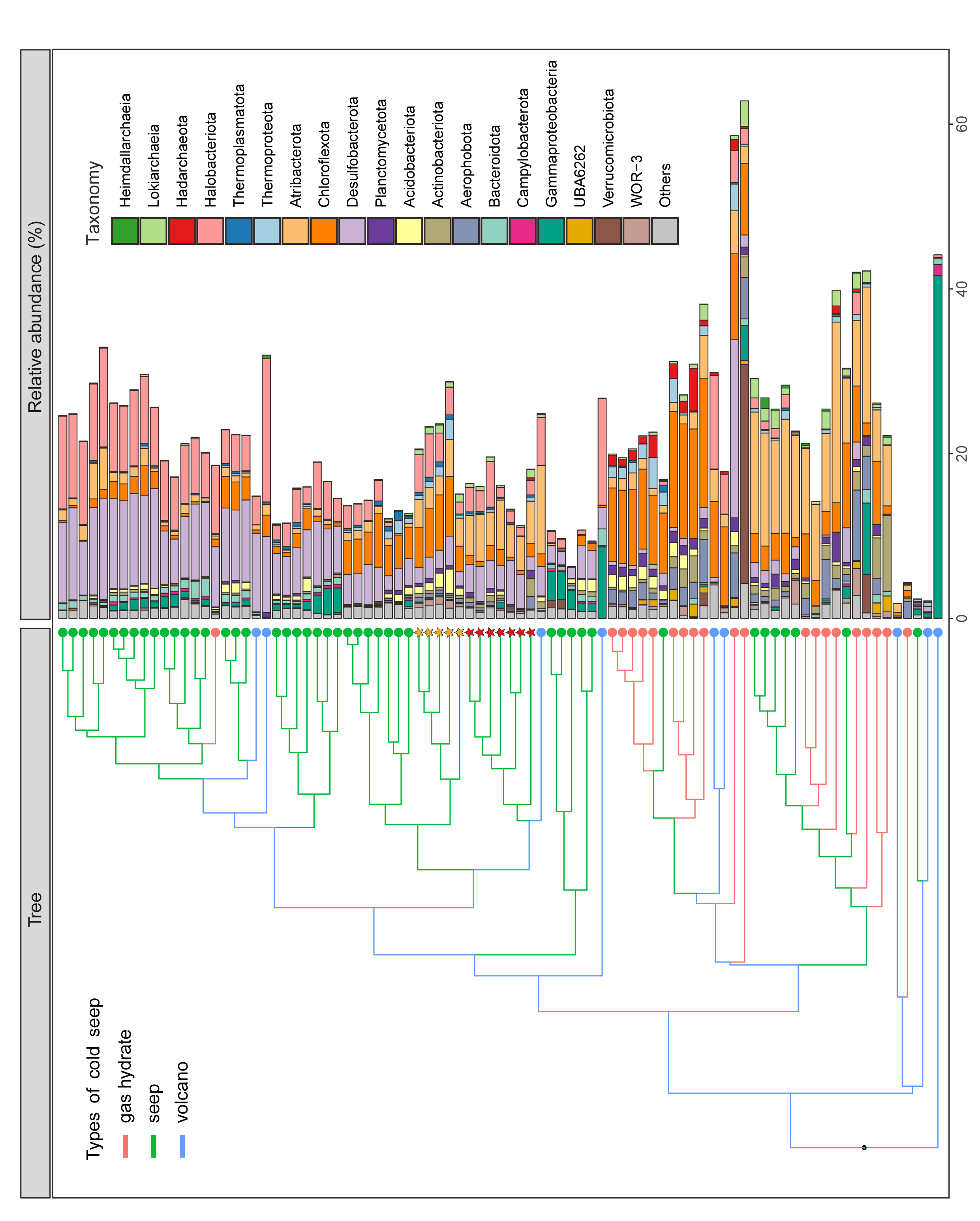
The community structures of microbiome involved in As cycling at cold seeps. The relative abundance of each MAG was estimated using CoverM. The compositions of microbiome involved in As cycling across different types of cold seep were clustered based the Bray–Curtis distance. Orange and red asterisks denote samples with lower and higher concentrations of Fe(II), respectively. Detailed statistics for As-related microbiome are provided in **Supplementary Data 4**.

### Expanded diversity of microbial lineages containing As detoxification genes

Among these As detoxification genes, *acr3, arsC1*/*arsC2, arsM* and *arsP* were widely distributed in bacteria and archaea, while other As detoxification genes (*arsB, arsI* and *arsH*) were sparsely distributed (**Figure 4**). The *acr3* gene is typically affiliated with Proteobacterial, Firmicutes, Actinobacterial and other bacterial sequences^36, 42, 46^. Our study observed an unexpectedly wider phylogenetical diversity of *acr3* than previously reported. Notably, Asgardarchaeota including Lokiarchaeia, Thorarchaeia, Sifarchaeia, LC30, along with Heimdallarchaeia and Wukongarchaeia described as the most likely sister group of eukaryotes, are firstly documented to have genetically capability for As(III) extrusion. The greater diversity of As detoxification genes found in Asgardarchaeota phylum further point to their ancient origin^19^. Furthermore, a considerable number of candidate bacterial phyla without cultured representatives (e.g. 4484-113, AABM5-125-24 and RBG-13-66-14) were also equipped with such an ability. Though their functional redundancy as As(III) efflux pumps, *arsB* was more phylogenetically conserved as compared with *acr3* and simply restricted to Alphaproteobacteria, Gammaproteobacteria and Campylobacterota (**Figure 4**). This observation is in agreement with previous reports comparing the diversity of *arsB* to *acr3*^36, 42, 46^. The *arsM* gene was relatively uncommon in terrestrial soil microorganisms^29, 36^. In contrast, this study showed that the *arsM* genes in seep microbes have a great taxonomic diversity similar to *acr3* gene, including Chloroflexota, Proteobacteria, Atribacterota, Asgardarchaeota, Hydrothermarchaeota, Thermoplasmatota, Thermoproteota as well as other currently unidentified bacterial phyla (e.g. 4484-113, AABM5-125-24 and RBG-13-66-14) (**Figure 4**). Among these, Atribacterota, Asgardarchaeota and the candidate bacterial phyla stated above have not previously been implicated in As methylation^19, 36^. For cytoplasmic As(V) reduction, Asgard archaeal lineages all lacked corresponding genes (*arsC1* and *arsC2*). The underlying causes to their absence in Asgardarchaeota are unclear. It’s likely that Asgardarchaeota lost cytoplasmic As(V) reduction genes during evolution or possess different enzyme systems. In general, these data advance our understanding on the phylogenetical diversity of As detoxification genes and highlight the potentially important role played by archaea in As cycling, Asgardarchaeota particularly.

**Figure 4.**
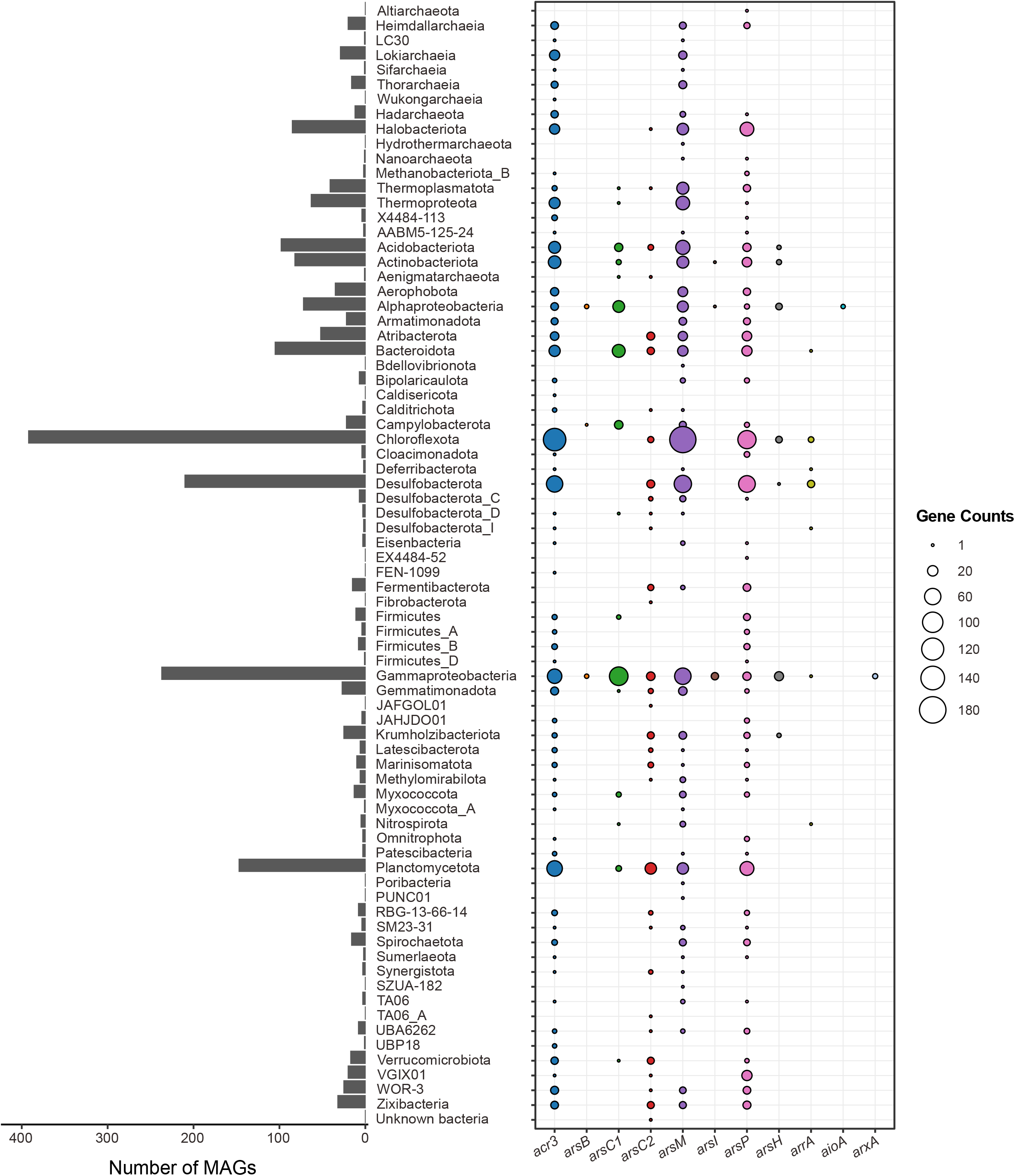
Phylogenetic distribution of As-cycling genes. Left bar plot showing the total number of genomes encoded in each phylogenetic cluster assigned by GTDB-Tk based on GTDB r207 release. Right bubble plot showing the number of As-cycling genes encoded within each phylogenetic cluster. Detailed information on phylogenetic diversity of As-cycling genes is provided in **Supplementary Data 5**.

### Diverse seep lineages are identified to perform As respiration

In addition to mitigating toxicity, some microorganisms can respire the redox-sensitive element of As to reap energetic gains (i.e. arsenotrophy), either via chemoautotrophic As(III) oxidation (*aioAB*/*arxAB*) or anaerobic As(V) respiration (*arrAB*)^47, 48^. The alpha subunits of these arsenotrophic enzymes form distinct clades with the dimethylsulfoxide (DMSO) reductase superfamily^34^. This superfamily also includes other enzymes critical in respiratory redox transformations, e.g. Nap and Nar. Here, we identified two AioA, three ArxA and 17 ArrA protein sequences, respectively. A phylogenetic analysis of recovered arsenotrophic protein sequences showed that they all clustered together with known AioA/ArxA and ArrA proteins (**Figure 5a**). Functional As bioenergetic *aioA*/*arxA* and *arrA* genes are generally found together with other necessary accessory genes. The *aioA* of As(III) oxidizing microorganisms always forms an operon with *aioB* and other genes involved in As detoxification and metabolisms (e.g. *aioD, aioXSR, arsR*)^15, 26^. Arx is demonstrated to be a variant of Arr and these two enzymes have a similar genetic arrangement. The *arrA*/*arxA* gene is always found together with the *arrB*/*arxB* and often with the *arrC*/*arxC* and *arrD*/*arxD*^15, 26^. The genomic organization analysis showed that identified two *aioA*, three *arxA* and 11 of 17 *arrA* genes all have corresponding accessory genes (**Figure 5b**), further confirming their potential identities as arsenotrophic enzymes.

**Figure 5.**
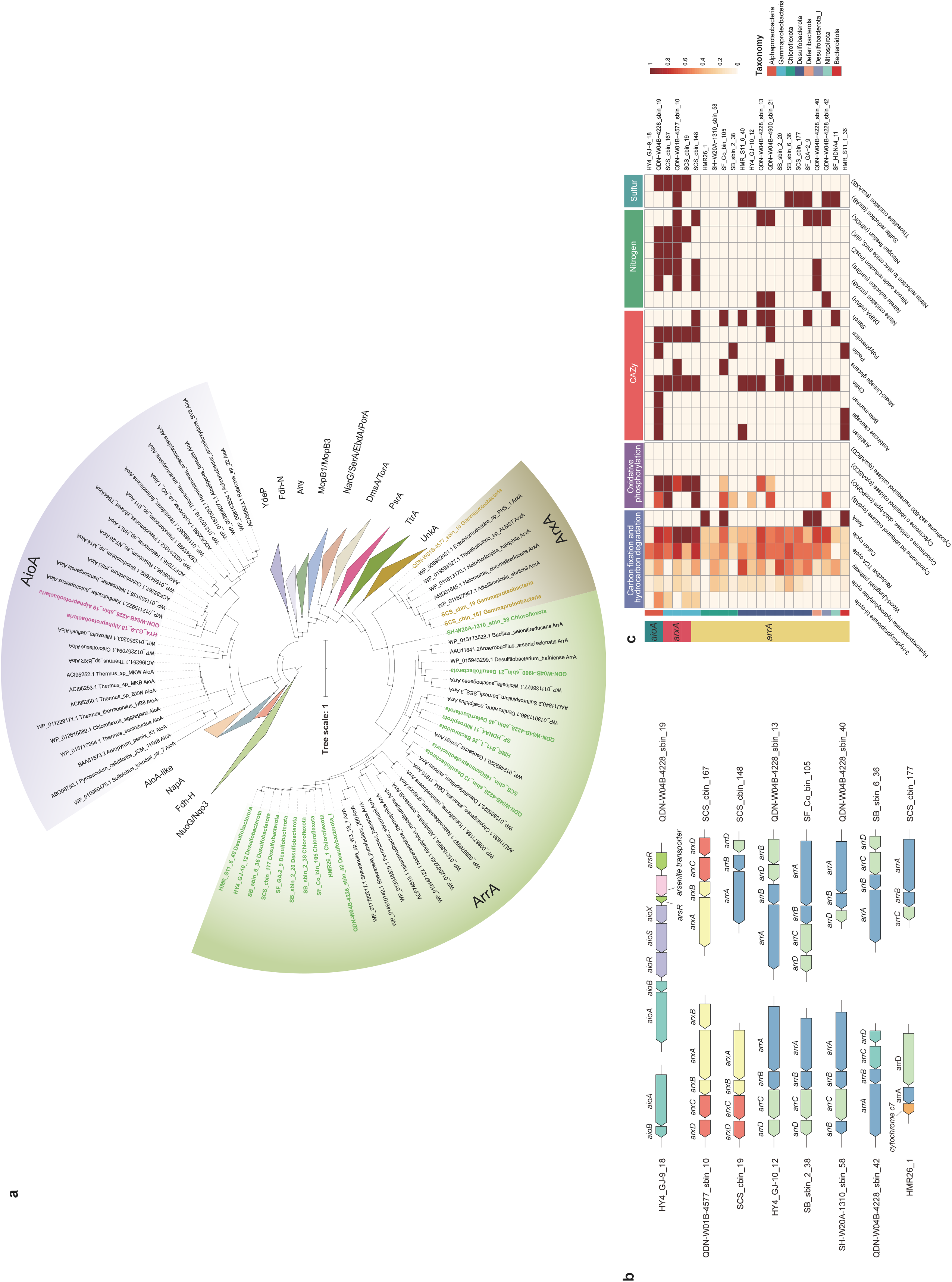
The metabolic potential of the arsenotrophic gene-carrying MAGs. (a) A maximum-likelihood tree of the DMSO reductase family, with protein sequences identified as associated with arsenotrophic enzymes in this study. Bootstrap values are generated from 1000 replicates. Bootstrap values≥70 are shown. Scale bar indicates amino acid substitutions per site (b) The genomic context of the *aioA, arxA* and *arrA* clusters in MAGs containing arsenotrophic genes. (c) Heatmap showing the predicted metabolism in potential As-respiring microbes. Detailed annotation is presented in **Supplementary Data 6**. The completeness of each pathway was calculated using the DRAM Distill function.

The *aioA*/*arxA* genes are uncommon in soil microbiomes and mostly found in Proteobacteria^15, 36, 49^. By assigning the taxonomy, *aioA*/*arxA* genes recovered here belonged to Gammaproteobacteria (*n* = 3) and Alphaproteobacteria (*n* = 2), consistent with previous findings (**Figure 3 and Figure 5c**). Nevertheless, 17 *arrA* genes were phylogenetically affiliated with seven distinct bacterial lineages: Bacteroidota (*n* = 1), Chloroflexota (*n* = 4), Deferribacterota (*n* = 1), Desulfobacterota (*n* = 8), Desulfobacterota_I (*n* = 1), Gammaproteobacteria (*n* = 1), and Nitrospirota (*n* = 1) (**Figure 3 and Figure 5c**). Despite that several other bacterial lineages (i.e. Deferribacterota, Firmicutes and Chrysiogenetes) are reported to contain *arrA* genes, most known As(V)-respiring microorganisms are assigned to proteobacterial clades^15, 36, 49^. Our findings, the *arrA*-containing Bacteroidota, Chloroflexota and Nitrospirota, expand the database of putative dissimilatory As(V) reducers.

### As respiration are potentially critical to central metabolisms in cold seeps

Microbially mediated As respiration has been verified to influence biogeochemical cycles of carbon and nitrogen, e.g. chemoautotrophic As(III) oxidation coupled with denitrification^50, 51^. Here, functional annotation identified near-complete calvin and reductive TCA carbon fixation pathways in *aioA*/*arxA*-carrying Alphaproteobacteria (*n =* 2) and Gammaproteobacteria MAGs (*n* = 3) (**Figure 5c**). Terminal reductase systems were also recognized in *aioA*/*arxA*-carrying MAGs, i.e. nitrate reductase (*narGHI*). The cooccurrence of these genes suggests that the As(III) oxidation may help support autotrophic carbon fixation and nitrate reduction.

In addition, five *arrA*-carrying MAGs possessed genes for AssA (**Figure 5c**), which mediate the first step of anaerobic activation of alkanes via fumarate addition^52^. Phylogenetic analyses revealed that identified AssA sequences were phylogenetically close to archaea-type and Group V AssA^53^ (**Supplementary Figure 5**). These potential hydrocarbon degraders were classified as Chloroflexota (*n* = 2), Deferribacterota (*n* = 1), Desulfobacterota (*n* = 1), and Bacteroidota (*n* = 1). Methane, the simplest hydrocarbon, has been demonstrated to stimulate As(V) respiration during the process of anaerobic oxidation of methane^54^. Similarly, the occurrence of both AssA and ArrA indicated that heterotrophic MAGs stated above may also employ As(V) as electron acceptor for anaerobic degradation of multi-carbon alkanes. Genes encoding carbohydrate-active enzymes (CAZymes) targeting various complex carbohydrates were also present in these arsenotrophic MAGs, including chitin, pectin, starch and plyphenolics (**Figure 5c**).

Notably, arsenotrophic MAGs may function as potential nitrogen fixers introducing new N to local environment. Genes encoding for the catalytic component of nitrogenase (i.e. *nifHDK*) were detected in one *arxA*-carrying (Gammaproteobacteria, *n* = 1) and six *arrA*-carrying (Gammaproteobacteria, *n* = 1; Desulfobacterota, *n* = 4; Desulfobacterota_I, *n* = 1) MAGs (**Figure 5c**). It has been previously reported that As(III) oxidation can fuel biological nitrogen fixation in tailing and metal(loid)-contaminated soils^55, 56^. The data present here further complement that diazotrophs could also fix N_2_ using energy obtained from dissimilatory As(V) reduction.

The metatranscriptomic reads were mapped against arsenotrophic MAGs to depict the gene expression profile at the genome level. Genes for dissimilatory As(V) reduction were transcriptionally active in the Jiaolong seep, as evidenced by the detection of *arrA* transcripts (21.7−340.1 TPM, **Supplementary Data 7**). In addition, transcripts of *assA* (9.2−6306.5 TPM) and *nifH* (155.1−28016.9 TPM) were identified in the Jiaolong seep, Shenhu area and Qiongdongnan Basin, implying anaerobic degradation of hydrocarbons and nitrogen fixation were actively expressed for these arsenotrophic microbes. No transcriptomic sequences related to *arrA, assA* and *nifH* genes were detected in the Haima seep. Nevertheless, it does not mean that genes of interest are not transcribed *in situ* because it is difficult to recover enough RNA from deep-sea samples and RNA can get lost during the process of deep-sea sampling^57^.

Our findings point towards a previously unrecognized arsenotrophs at seeps, impacting both carbon and nitrogen cycling. However, we acknowledge that cultivation experiments with As-respiring isolates are ultimately needed both to elucidate their lifestyle and confirm functionality for As-dependent carbon fixation, hydrocarbon and carbohydrate degradation as well as nitrogen fixation.

## Conclusions

Microbial transformation of As has been well documented and characterized in environments in such as ocean water, groundwater and geothermal springs, but the knowledge on gene- and genome-level As cycling in deep sea (e.g. cold seep) is limited. Our study demonstrated that As methylation and cytoplasmic As(V) reduction were the predominant detoxification mechanisms employed by cold seep microbiomes. These results substantially expanded the diversity of As detoxification genes to a broader microbial community including Asgardarchaeota and a great number of candidate bacterial phyla. In addition, diverse arsenotrophic lineages are also identified, including Bacteroidota, Chloroflexota, Nitrospirota, etc, which also potentially participate in carbon and nitrogen biogeochemical cycling. This study provides a detailed understanding of As biotransformation in a complex microbiome in deep-sea realms, which could have significant implications for addressing environmental issues. Our result will also provide insights for microbial evolution in the early ocean with harmful metal(loids), e.g. As, as a driving force^58^.

## Methods

### Metagenomic and metatranscriptomic data sets

The 87 metagenomes and 33 metatranscriptomes analyzed in this study are derived from 13 globally distributed cold seep sites (**Supplementary Figure 1**). Among them, 65 metagenomes and 10 metatranscriptomes were compiled from our previous publications^8, 59^, and other 22 metagenomes were downloaded from NCBI Sequencing Read Archive (SRA). A detailed description of sampling locations and sequencing information for metagenomic and metatranscriptomic data is given in **Supplementary Data 1**.

### Bioinformatic analyses

DNA reads pre-processing, metagenomic assembly and binning were performed with the function modules of metaWRAP (v1.3.2)^60^. First, the metaWRAP Read_qc module was used to trim raw sequencing DNA reads. Then the filtered DNA reads were individually assembled with the metaWRAP Assembly module using Megahit^61^ or metaSPAdes^62^ with default settings (detailed assembly statistics are summarized in **Supplementary Data 1**). In addition, metagenomic reads from the same sampling station (*n* = 10) were also co-assembled using Megahit with the default settings. Thereafter, MAGs were recovered from contigs with the length longer than 1kb using the metaWRAP Binning module (parameters: -maxbin2 -concoct -metabat2) or the VAMB tool^63^ (v3.0.1; default parameters; detailed binning statistics are summarized in **Supplementary Data 1**). Further refinement of MAGs was performed by the Bin_refinement module of metaWRAP (parameters: -c 50 -x 10), and CheckM (v1.0.12)^64^ was used to estimate the completeness and contamination of these MAGs. All MAGs were dereplicated at 95% average nucleotide identity (ANI) using dRep (v3.4.0; parameters: -comp 50 -con 10)^65^ to obtain representative species MAGs. This analysis provided a non-redundant genome set consisting of 1,741 species-level MAGs.

Raw metatranscriptomes were quality filtered with the Read_qc module of metaWRAP (v1.3.2)^60^ as described above. The removal of ribosomal RNAs was conducted with sortmeRNA (v2.1)^66^ in the quality-controlled metatranscriptomic reads.

### Non-redundant gene catalog construction

Genes were predicted on contigs (≥1kb) from the assemblies using the METABOLIC pipeline (v4.0)^67^, which resulted in 33,799,667 protein-coding genes. Clustering of the predicted proteins was performed with MMseqs2 (v13.45111)^68^ using the cascaded clustering algorithm at 95% sequence similarity and 90% sequence coverage (parameters: -c 0.95 -min-seq-id 0.95 -cov-mode 1 -cluster-mode 2) following the ref.^69^. This process yielded a total of 17,217,131 non-redundant gene clusters.

### Searching for As cycling genes

In this study, 11 well-characterized marker genes^70, 71^ were selected to assess their potential influence to the As biogeochemical cycle. These genes include eight As detoxification genes (*acr3, arsB, arsC1, arsC2, arsP, arsH, arsI*, and *arsM*) and three As respiratory genes (*aioA, arrA*, and *arxA*). A hidden Markov model (HMM)-based search was performed to identify As-related genes in non-redundant gene catalogue by using hmmsearch function in HMMER package (v3.1b2)^72^. The HMM profile searches and score cutoffs for 11 As-related genes were taken from Lavy et al. (2020)^71^.

### Taxonomic and functional profiling of MAGs

As-related MAGs were taxonomically annotated using the classify_wf function of the GTDB-Tk toolkit (v2.1.1)^73^ with default parameters against the GTDB r207 release. For all MAGs, gene calling and metabolic pathway prediction were conducted with the METABOLIC pipeline (v4.0)^67^. Functional annotation of genomes was also carried out by searching against KEGG, Pfam, MEROPS and dbCAN databases using DRAM (v1.3.5)^74^. The identification of As-related genes in MAGs was performed by searching against As-related HMM profiles from Lavy et al. (2020)^71^ as reported above. Genes involved in anaerobic hydrocarbon degradation were screened using BLASTp (identity >30%, coverage >90%, e < 1 × 10^−20^) against local protein databases^53^.

### Abundance calculations

For contig level, the relative abundance of genes related to As cycling across 87 metagenomes were calculated from non-redundant gene catalog using the program Salmon (v1.9.0)^75^ in the mapping-based mode (parameters: -validateMappings -meta). GPM (genes per million) values were used as a proxy for gene abundance as describe in ref.^74^. For genome level, the relative abundance of each MAG was profiled by mapping quality-trimmed reads from the 87 metagenomes against the MAGs using CoverM in genome mode (https://github.com/wwood/CoverM) (v0.6.1; parameters: -min-read-percent-identity 0.95 -min-read-aligned-percent 0.75 -trim-min 0.10 -trim-max 0.90 -m relative_abundance).

To calculate the transcript abundances of As-related genes, we also mapped clean reads from the 33 metatranscriptomes to non-redundant gene catalog or arsenotrophic MAGs. The transcript abundance of each gene was calculated as the metric-TPM (transcripts per million). GPM or TPM values were normalized based on the gene length and sequencing depth.

### Phylogenetic analyses of functional genes

For phylogeny inference, protein sequences of functional genes were aligned with MAFFT (v7.490, -auto option)^76^, and gap sequences were trimmed using trimAl (v. 1.2.59, -automated1 option)^77^. Maximum likelihood phylogenetic trees were constructed for each genes using IQ-TREE (v2.12)^78^ with the following options: -m TEST -bb 1000 -alrt 1000. Branch support was estimated using 1000 replicates of both ultrafast bootstrap approximation (UFBoot)) and Shimodaira-Hasegawa (SH)-like approximation likelihood ratio (aLRT). Reference protein sequences for As-based respiratory cycle were obtained from Saunders et al. (2019)^34^. Reference protein sequences for fumarate addition were derived from Zhang et al. (2021)^53^. All the tree files were uploaded to Interactive tree of life (iTOL; v6)^79^ for visualization and annotation.

### Statistical analyses

Statistical analyses were done in R (v4.0.4-v4.1.0) with the following descriptions. Normality and homoscedasticity of data were evaluated using Shapiro-Wilk test and Levene’s test, respectively. One-way analysis of variance (ANOVA) and least significant difference (LSD) test were conducted to evaluate the variations of each gene across different sediment depths and types of cold seeps. The partial least squares discrimination analysis (PLS-DA) was performed based on the GPM values of As-cycling genes with R package ‘mixOmics’. The permutational multivariate analysis of variance (PERMANOVA) was employed to test whether As-cycling genes shifted among different sediment depths and types of cold seeps using ‘adnois’ function in vegan package. All PERMANOVA tests were performed with 9999 permutations based on Bray–Curtis dissimilarity.

## DATA AVAILABILITY

Non-redundant gene catalog, assemblies, MAGs containing As cycling genes, and raw tree files have been uploaded to Figshare (https://figshare.com/s/833c3dc27319617e76ed). Arsenotrophic MAGs have also been deposited in NCBI under accession numbers SAMN33581604-33581625 (BioProject ID PRJNA831433).

## CODE AVAILABILITY

The present study did not generate codes, and mentioned tools used for the data analysis were applied with default parameters unless specified otherwise.

## ACKNOWLEDGMENTS

This study was supported by the Scientific Research Foundation of Third Institute of Oceanography, MNR (No. 2022025), State Key Laboratory of Marine Geology, Tongji University (No. MGK202303), China Postdoctoral Science Foundation (2022M723709), the Science and Technology Projects in Guangzhou (No. 202102020970), the Guangdong Basic and Applied Basic Research Foundation (No. 2019B030302004, 20201910240000691) and the Marine Geological Survey Program of China Geological Survey (DD20221706).

## COMPETING INTERESTS

The authors declare no competing interests.

## AUTHOR CONTRIBUTIONS

XD designed this study with input from JL. CZ and XL analyzed omic data. XD, CZ, XL, LDS and ZS interpreted data. JL and XX contributed to data collection. XD and CZ drafted the paper, with input from all other authors.

## Notes

### Competing Interest Statement

The authors have declared no competing interest.

### Summary of Updates

The title was revised.

