## Supplementary Information for "Unexpected genetic and microbial diversity for arsenic cycling in deep sea cold seep sediments"

### Supporting Information

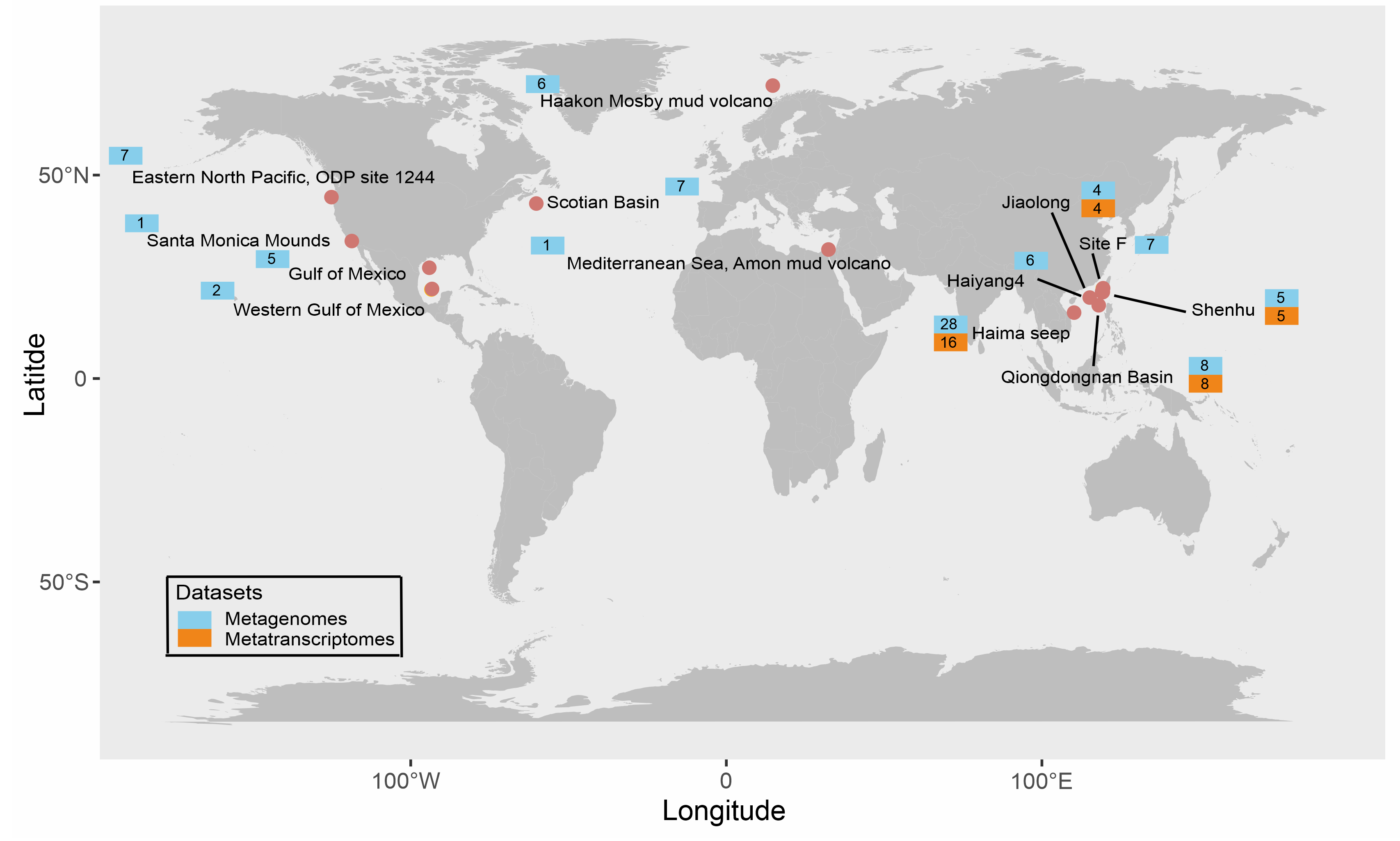

**Figure S1**. Geographical distributions of 13 cold seep sites analyzed in this study. Colors indicate the type of samples collected at each site: metagenome (blue); metatranscriptome (orange). The numbers indicate the number of samples for metagenomes or metatranscriptomes. Further details for each metagenome and metatranscriptome can be found in **Table S1**.

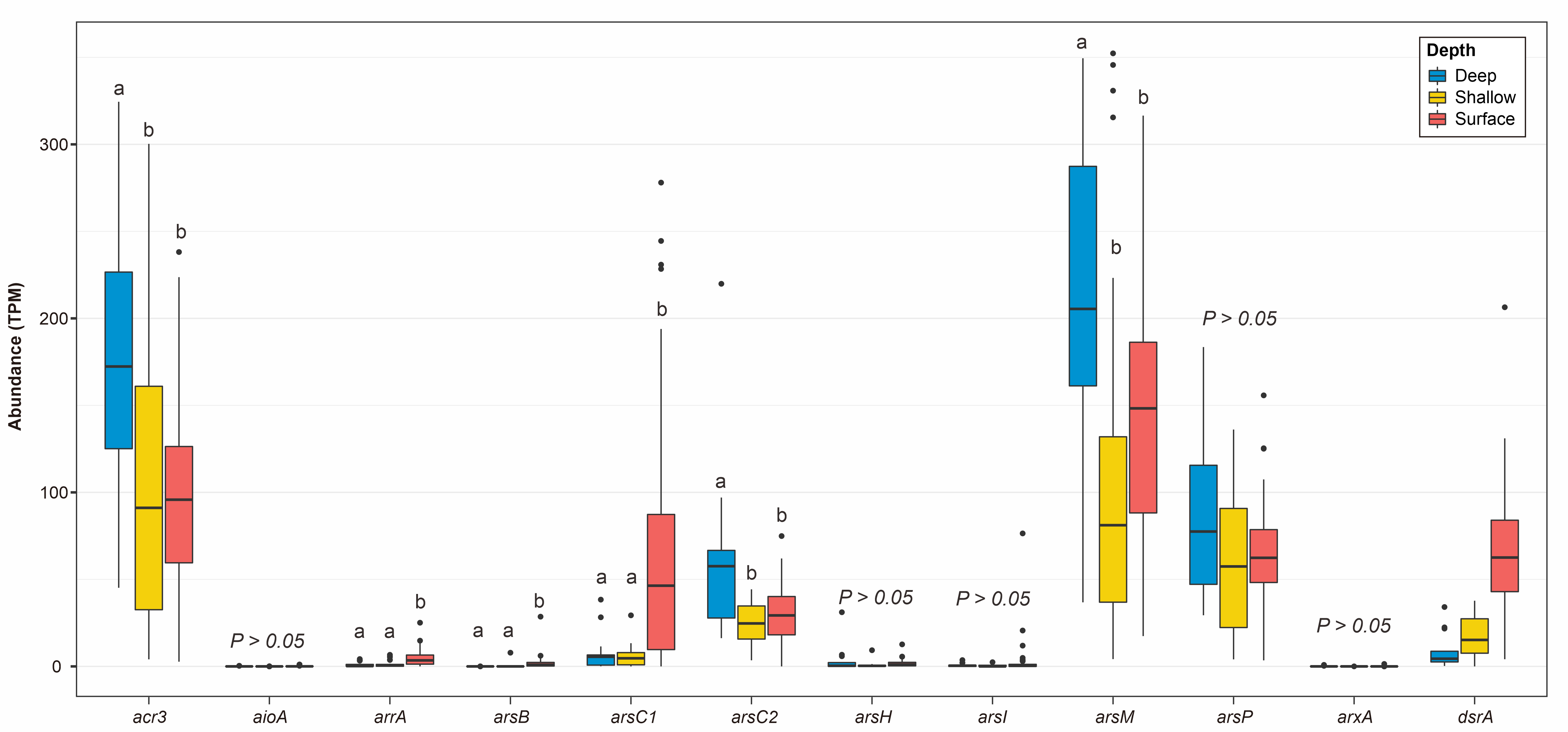
**Figure S2**. The abundances of As-cycling gene across different sediment depths with *drsA* gene as a comparison.

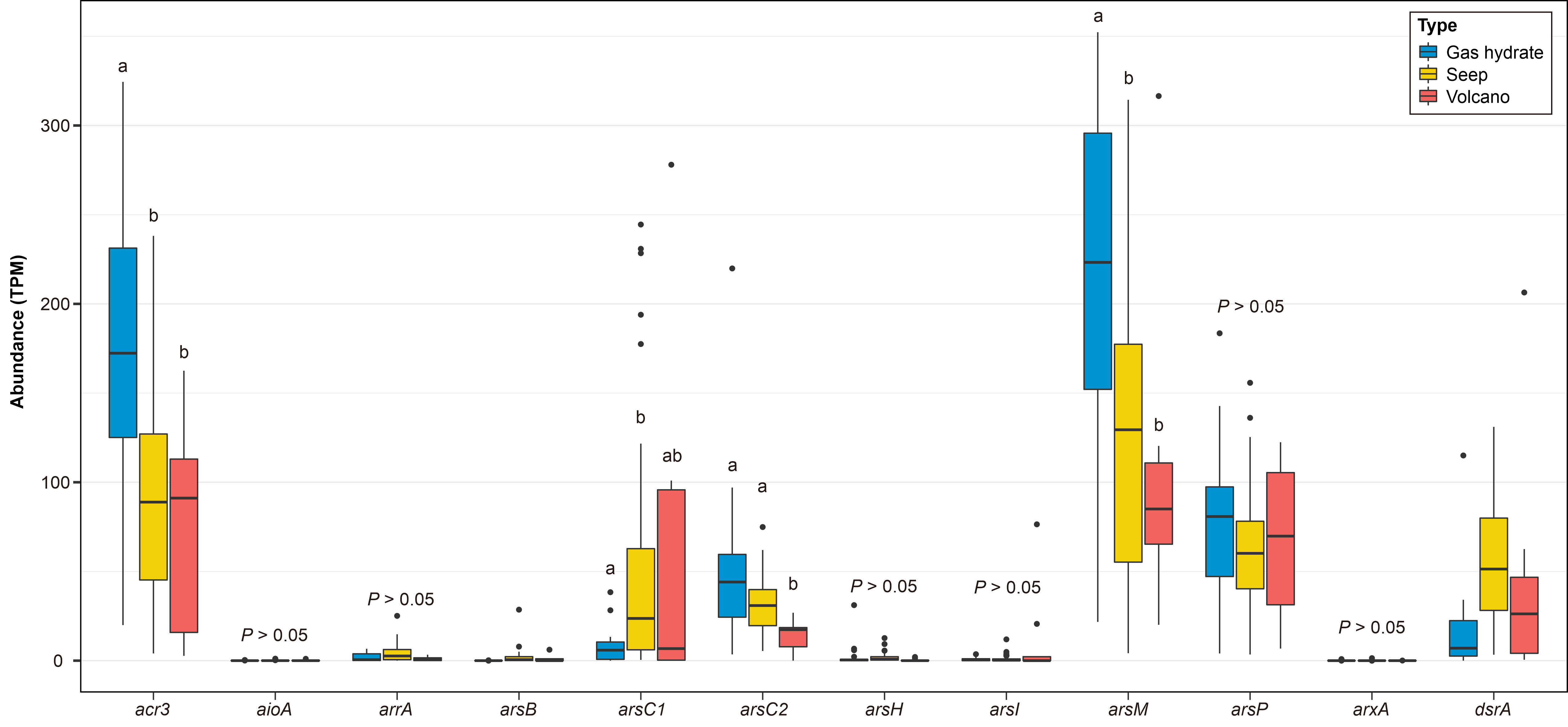

**Figure S3**. The abundances of As-cycling gene across different types of cold seep ecosystems with *drsA* gene as a comparison.

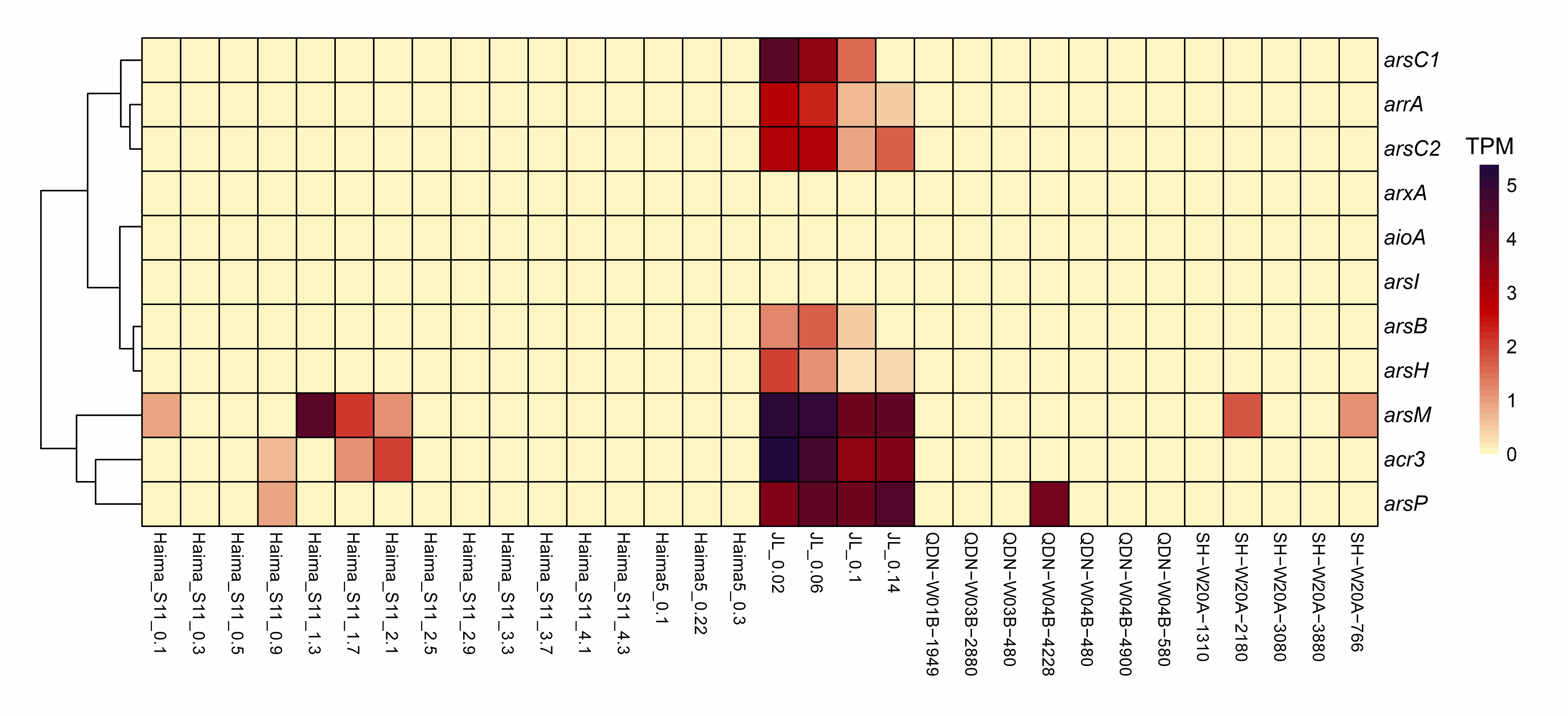

**Figure S4**. The abundance of As-cycling transcripts in 33 sediment metatranscriptomes.

**Figure S5**. Phylogenetic analysis of identified AssA protein sequences in *arrA*-carrying MAGs. Bootstrap values >70% are shown as black dots on branch nodes. Scale bar represents amino acid substitutions per site.
